# The evolutionary impact of Intragenic FliA Promoters in Proteobacteria

**DOI:** 10.1101/188474

**Authors:** Devon M. Fitzgerald, Carol Smith, Pascal Lapierre, Joseph T. Wade

## Abstract

Recent work has revealed that large numbers of promoters in bacteria are located inside genes. In contrast, almost all studies of transcription have focused on promoters upstream of genes. Bacterial promoters are recognized by Sigma factors that associate with initiating RNA polymerase. In *Escherichia coli*, one Sigma factor recognizes the majority of promoters, and six “alternative” Sigma factors recognize specific subsets of promoters. One of these alternative Sigma factors, FliA (σ^28^), recognizes promoters upstream of many flagellar genes. We previously showed that most *E. coli* FliA binding sites are located inside genes. However, it was unclear whether these intragenic binding sites represent active promoters. Here, we construct and assay transcriptional promoter-*lacZ* fusions for all 52 putative FliA promoters previously identified by ChIP-seq. These experiments, coupled with integrative analysis of published genome-scale transcriptional datasets, reveal that most intragenic FliA binding sites are active promoters that transcribe highly unstable RNAs. Additionally, we show that widespread intragenic FliA-dependent transcription is a conserved phenomenon, but that the specific promoters are not themselves conserved. We conclude that intragenic FliA-dependent promoters and the resulting RNAs are unlikely to have important regulatory functions. Nonetheless, one intragenic FliA promoter is broadly conserved, and constrains evolution of the overlapping protein-coding gene. Thus, our data indicate that intragenic regulatory elements can influence protein evolution in bacteria, and suggest that the impact of intragenic regulatory sequences on genome evolution should be considered more broadly.

**AUTHOR SUMMARY:** Recent genome-scale studies of bacterial transcription have revealed thousands of promoters inside genes. In a few cases, these promoters have been shown to transcribe functional RNAs. However, it is unclear whether most intragenic promoters have important biological function. Similarly, there are likely to be thousands of intragenic binding sites for transcription factors, but very few have been functionally characterized. Moreover, it is unclear what impact intragenic promoters and transcription factor binding sites have on evolution of the overlapping genes. In this study, we focus on FliA, a broadly conserved Sigma factor that is responsible for initiating transcription of many flagellar genes. We previously showed that FliA directs RNA polymerase to ~50 genomic sites in *Escherichia coli*. In our current study, we show that while most intragenic FliA promoters are actively transcribed, very few are conserved in other species. This suggests that most FliA promoters are not functional. Nonetheless, one intragenic FliA promoter is highly conserved, and we show that this promoter constrains evolution of the overlapping protein-coding gene. Given the enormous number of regulatory DNA sites within genes, we propose that the evolution of many genes is constrained by these elements.

## INTRODUCTION

In bacteria, RNA polymerase (RNAP) requires a transcription initiation factor, σ, to recognize promoter elements and initiate transcription. Bacteria encode one housekeeping σ factor that functions at most promoters, and multiple “alternative” σ factors that each recognize smaller sets of promoters. Historically, promoters were thought to be located solely upstream of annotated genes. However, widespread transcription initiation from inside genes has now been described in *Escherichia coli* and many other species (reviewed, [1,2]). Consistent with these observations, the *E. coli* housekeeping σ factor, σ^70^, has been shown to bind many intragenic sites [3]. Similar findings have been reported for alternative σ factors, e.g. 40% of *Mycobacterium tuberculosis* SigF binding sites, 25% of *E. coli* σ^32^ binding sites, and 62% of *E. coli* σ^54^ binding sites are inside genes [4-7]. The high degree of pervasive transcription involving multiple σ factors suggests that intragenic promoters have a substantial impact on global transcriptional networks.

Like σ factors, DNA-binding transcription factors often bind extensively within genes [2,7-11]. The regulons of most transcription factors have not been mapped, even for *E. coli*, suggesting that most intragenic binding sites remain to be identified. Indeed, a study of 51 transcription factors in *Mycobacterium tuberculosis* suggests that a typical bacterial genomes contain >10,000 intragenic binding sites [10]. The transcriptional activity of most intragenic transcription/σ factor binding sites have not been extensively studied, but many are likely to be functional [10]. Although transcription regulatory networks evolve rapidly, individual regulatory interactions are often subject to positive selective pressure [12-14]. Hence, many intragenic transcription/σ factor binding sites are maybe functional, and hence are likely to be conserved. A previous study suggested that positive selection on intragenic transcription/σ factor binding sites in human cells constrains the evolution of overlapping protein-coding genes [15]. The impact of bacterial intragenic binding sites on overlapping protein-coding genes has not been assessed.

FliA (σ^28^) is an alternative σ factor involved in transcription of genes associated with flagellar motility and chemotaxis (reviewed [16]). FliA also initiates transcription of some non-flagellar genes in *E. coli* [17], and is encoded by some non-motile bacteria, such as *Chlamydia* [18], suggesting additional non-flagellar roles. Recently, we reported that over half of *E. coli* FliA binding sites are located inside genes, often far from gene starts [17]. These intragenic sites were split approximately evenly between those occurring in the sense and antisense orientations, with respect to the overlapping gene. Most intragenic FliA binding sites were not associated with detectable FliA-dependent RNAs, so it is unclear whether they represent functional promoters. Notably, FliA is the most highly and broadly conserved alternative σ factor [16,19]. The interactions between FliA, RNA polymerase, and promoter DNA are so highly conserved that the *Bacillus subtilis* homolog, σ^D^, can complement an *E. coli* Δ*fliA* strain [20]. Like many alternative σ factors, FliA has a decreased ability to melt DNA as compared to housekeeping σ factors [19,21]. Thus, FliA-dependent transcription initiation requires a stringent match to its consensus promoter sequence [22]. Together, the high conservation and readily identifiable motif make FliA a good model for evolutionary analysis of intragenic σ factor binding.

In this study, we evaluate the promoter activity of intragenic FliA binding sites in *E. coli.* We also assess the conservation of intragenic FliA promoters and map the *Salmonella* FliA regulon. We conclude that most intragenic FliA binding sites represent bona fide promoters that transcribe unstable intragenic RNAs. We show that extensive intragenic transcription by FliA is a conserved phenomenon, but the genetic locations of promoters are generally not conserved. Nonetheless, we show that a single intragenic FliA promoter is under strong selective pressure that constrains the evolution of the FlhC protein. This is the first documented example of intragenic regulatory sequence impacting evolution of the overlapping protein-coding gene, and suggests that selective pressure on intragenic binding sites for σ factors and transcription factors is an overlooked factor in protein evolution in compact bacterial genomes.

## RESULTS

### Most intragenic FliA binding sites represent transcriptionally active promoters

To test whether FliA binding sites previously identified by ChIP-seq [17] represent active promoters, we generated transcriptional fusions of potential promoters to the *lacZ* reporter gene. For each of the 52 putative FliA promoters, the region from approximately −200 to +10 was cloned upstream of *lacZ* on a single-copy plasmid (Figure 1 A). We chose to include 200 bp upstream sequence because at least one FliA promoter is regulated by a transcription factor binding upstream [23]. Plasmids were transformed into a motile strain of *E. coli* MG1655 (i.e. expressing FliA), or an isogenic Δ*fliA* derivative, and assayed for β-galactosidase activity. Of the 20 intergenic promoters, 15 displayed significant FliA-dependent activity (*t*-test, *p* ≤0.05; Figure 1B). Of the 30 intragenic promoters, 10 out of 16 sense-and 7 out of 14 antisense-orientation putative intragenic promoters showed significant FliA-dependent activity (*t*-test, *p* ≤0.05; Figure 1C). These intragenic FliA-dependent promoters include all five that have been previously associated with transcription of stable RNAs *((flhC*)*motAB-cheAW, (yafY)ykfB, (yjdA)yjcZ, (uhpT)*, and antisense *(hypD)*, where genes in parentheses indicate those with an internal FliA promoter). One of the two putative promoters located in convergent intergenic regions also showed significant FliA-dependent activity (*t*-test, *p* ≤0.05; Figure 1C). It should be noted that some fusions had very high levels of background activity, which may have prevented the detection of lower levels of FliA-dependent transcription from these promoter fusions. Of note, no FliA-dependent activity was detected for the well-characterized promoters upstream *of fliA, fliD*, an*d fliL*, likely due to overwhelming transcriptional activity from the strong, σ^70^-dependent, FlhDC-activated promoters known to be immediately upstream [17,24,25]. High β-galactosidase activity associated with the *lacZ* fusions for *pntA, cvrA, glyA, proK*, and *insB-4/cspH* suggest they are also likely to include σ^70^ promoters that may preclude identification of FliA-dependent transcription. Consistent with this, we previously detected σ^70^ binding sites <200 bp upstream of all of these putative FliA promoters [3].

**Figure 1.**
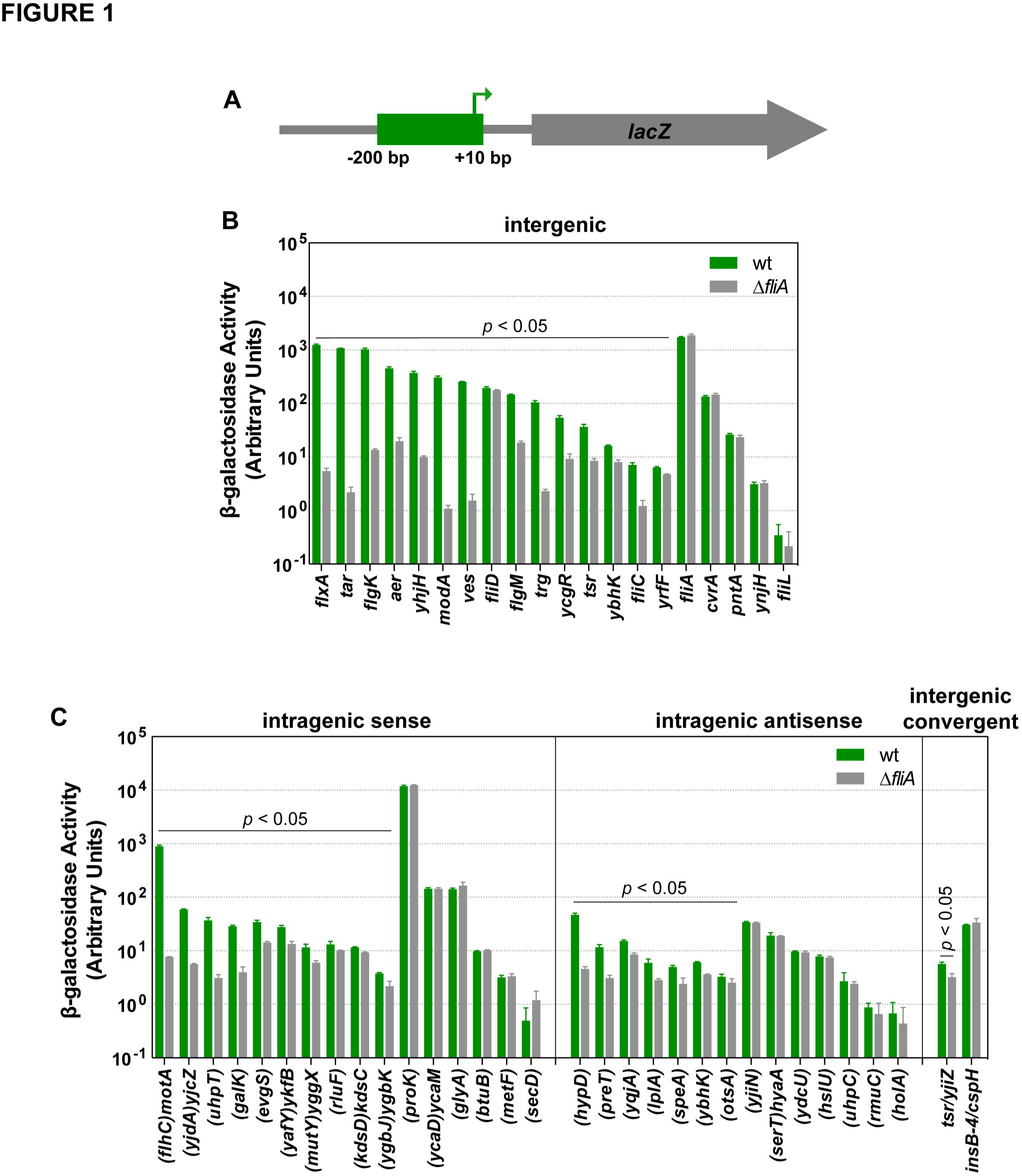
Identification of transcriptionally active FliA binding sites using reporter gene fusions. **(A)** Schematic of transcriptional fusions of potential FliA promoters to the *lacZ* reporter gene. For all FliA binding sites identified in a previous study, transcriptional fusions to *lacZ were* constructed using positions −200 to +10 relative to the predicted TSS based on the previously identified FliA binding motif [17]. **(B)** β-galactosidase activity for transcriptional fusions for FliA binding sites in intergenic regions upstream of genes, for wild-type (wt; DMF122; green bars) and Δ*fliA* (DMF123; gray bars) cells. Reporter fusions that showed significantly lower β-galactosidase activity in Δ*fliA* cells than wild-type cells (*t*-test *p* < 0.05) are indicated. The genes downstream of the FliA binding sites are listed on the x-axis. **(C)** As above, but for FliA binding sites within genes or between convergently transcribed genes. Genes containing FliA binding sites are listed on the x-axis in parentheses. Genes not in parentheses are downstream of the corresponding FliA binding site. Error bars indicate one standard deviation from the mean (n = 3).

We previously identified FliA-regulated transcripts using RNA-seq, although most intragenic FliA sites were not associated with a detectable RNA [17]. However, this method often fails to detect unstable RNAs. To independently assess whether intragenic FliA binding sites act as promoters, we analyzed two published datasets generated from motile *E. coli* strains: (i) genome-wide transcription start site (TSS) mapping by differential RNA-seq (dRNA-seq) [26], and (ii) Nascent Elongating Transcript sequencing (NET-seq) [27]. dRNA-seq identifies TSSs by selectively degrading processed transcripts bearing a 5’ monophosphate, and then preparing a library of the remaining 5’ triphosphate-bearing primary transcripts [28]. By focusing reads to the 5’ ends of transcripts, this technique is more sensitive than standard RNA-seq, and can distinguish intragenic RNAs from overlapping mRNAs. NET-seq isolates nascent RNA still bound to RNAP, facilitating detection of unstable transcripts prior to degradation [29].

To compare FliA binding site location to TSS mapping data, we determined the distance from the predicted FliA promoter sequence associated with each FliA binding site [17] to all downstream TSSs within 500 bp (Figure 2A). For most well-characterized FliA-dependent promoters for flagellar genes, the distance between the center of the promoter sequence and TSS was between 18 and 22 bp. For other FliA binding sites, we observed a strong enrichment for TSSs between 18 and 23 bp downstream of FliA motif centers. In total, 38 of the 52 FliA binding sites have a TSS located 18-23 bp downstream of the center of their predicted promoter. This positional enrichment is highly significant when compared to the same analysis performed with a randomized TSS dataset; only one random TSS was between 18-23 bp downstream of a FliA motif center (Fisher’s exact test, p<0.0001).

**Figure 2.**
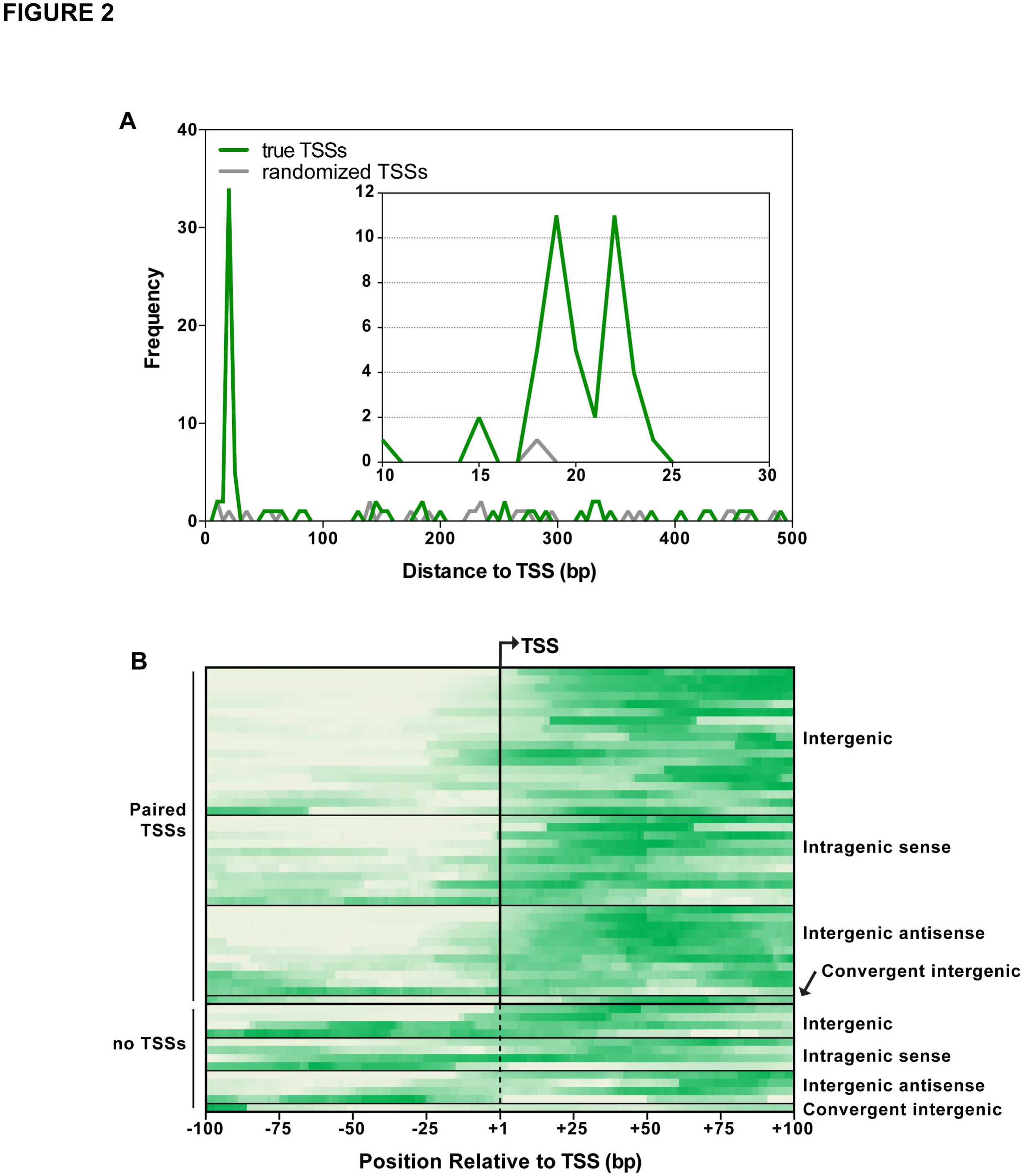
Identification of transcriptionally active FliA binding sites by mining genome-scale transcriptome datasets. **(A)** For each FliA binding site identified previously [17], we determined the distance to each downstream TSS identified by [26] within a 500 bp range. The frequencies of these distances are plotted in 10 bp bins (green line), with the inset showing the frequency of binding sites 10-30 bp upstream of TSSs with a bin size of 1 bp. The gray line shows the frequency of distances from FliA binding sites to a control, randomized TSS dataset (see Methods). **(B)** Normalized sequence read coverage from published NET-seq data [27] (see Methods) for each previously identified FliA binding site [17], plotted 100 bp upstream and downstream of the known/predicted TSS. Predicted TSSs are indicated by the dashed vertical line. Darker green indicates higher sequence read density.

To systematically assess whether FliA binding sites are associated with signal in the NET-seq dataset, the sequence read coverage upstream and downstream of FliA binding sites was determined. For FliA binding sites associated with a TSS, the read coverage at each position from −100 to +100 was determined relative to the TSS. For all other FliA binding sites, a TSS was predicted at 20 bp downstream of the predicted promoter sequence center (average position of other TSSs), and coverage was determined from −100 to +100 relative to this position. The coverage profile for each binding site was normalized to the minimum and maximum coverage in the region and plotted as a heatmap (Figure 2B). There is a clear trend of higher NET-seq read coverage downstream of FliA binding sites, compared to the regions immediately upstream. To quantify this trend, the ratio of NET-seq read coverage upstream and downstream of the TSS was calculated for each putative FliA-dependent promoter. In total, 44 out of the 52 putative promoters showed at least 2-fold higher coverage in the region 100 bp downstream of the TSS than in the region 100 bp upstream of the TSS. These 44 putative promoters included 19 that are intragenic (Table 1). As expected, there is a high degree of overlap between the FliA binding sites with transcriptional activity detected by NET-seq and those detected by TSS association (Table 1).

**Table 1.**
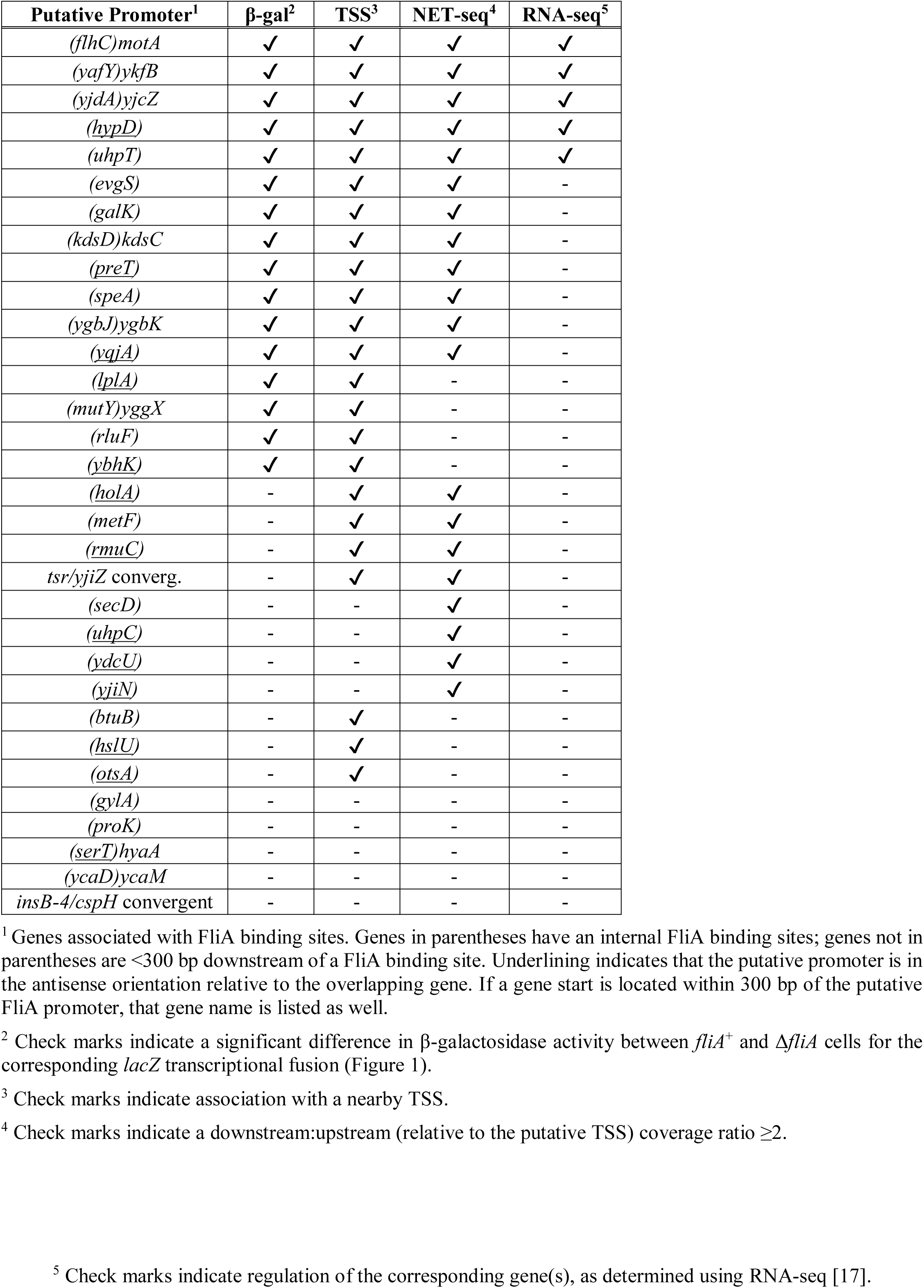
Intragenic FliA binding sites show evidence of transcriptional activity.

In total, 26 of the 30 intragenic FliA binding sites, and one of the two FliA sites in a convergent intergenic region, show evidence of promoter activity from at least one assay. Table 1 summarizes the existing evidence for these sites. It should be noted that neither the TSS nor NET-seq datasets have matched Δ*fliA* controls, so it is formally possible that TSSs/transcripts are associated with FliA-independent promoters. However, this is highly unlikely given the position of putative TSSs and the position of NET-seq signal with respect to the predicted FliA promoter sequences. Overall, there is substantial overlap between the sets of putative intragenic promoters that display FliA-dependent activity in promoter fusion assays, those with appropriately positioned TSSs, and those that have high NET-seq read coverage ratios (downstream: upstream).

### Intragenic FliA promoters are not conserved

To assess whether intragenic FliA promoters and binding sites are likely to be functionally important, we determined conservation of these sites bioinformatically. The sequence surrounding each of the 52 FliA binding sites previously identified by ChIP-seq [17] was extracted and used as a BLAST query to search genomes from 24 γ-proteobacterial genera. All genomes queried encode FliA, except for those *of Klebsiella* and *Raoultella*, which were included as controls. If a homologous region was identified, it was scored against the previously determined *E. coli* FliA position-weight matrix [17]. These scores are depicted as a heatmap in Figure 3, where yellow represents the highest-scoring sites and blue the lowest-scoring. Sites are ranked by total degree of conservation, from top to bottom. The well-characterized FliA-dependent promoter inside *flhC*, which drives transcription of the downstream *motABcheAW* operon, was the most highly conserved. All other well-characterized, flagellar-related FliA promoters were well-conserved at the sequence level, with the exception of the promoter upstream of the *fliLMNOPQR* operon, which is also transcribed by σ^70^ in *E. coli.* Most novel intergenic and intragenic FliA binding sites showed no evidence of conservation, even in close relatives such as *Salmonella.* It should be noted that a few intragenic FliA binding sites, such as those inside *hslU, glyA*, and *ybhK*, appear conserved, but score equally well in species that *lack fliA (Klebsiella* and *Raoultella)*, suggesting they are maintained for reasons independent of their ability to bind FliA, most likely because of high levels of conservation for these protein-coding genes. A few other intragenic promoters, such as those inside *uhpC, hypD, metF*, and *speA*, show possible sequence conservation in *Salmonella*, but not in more distantly related genera.

**Figure 3.**
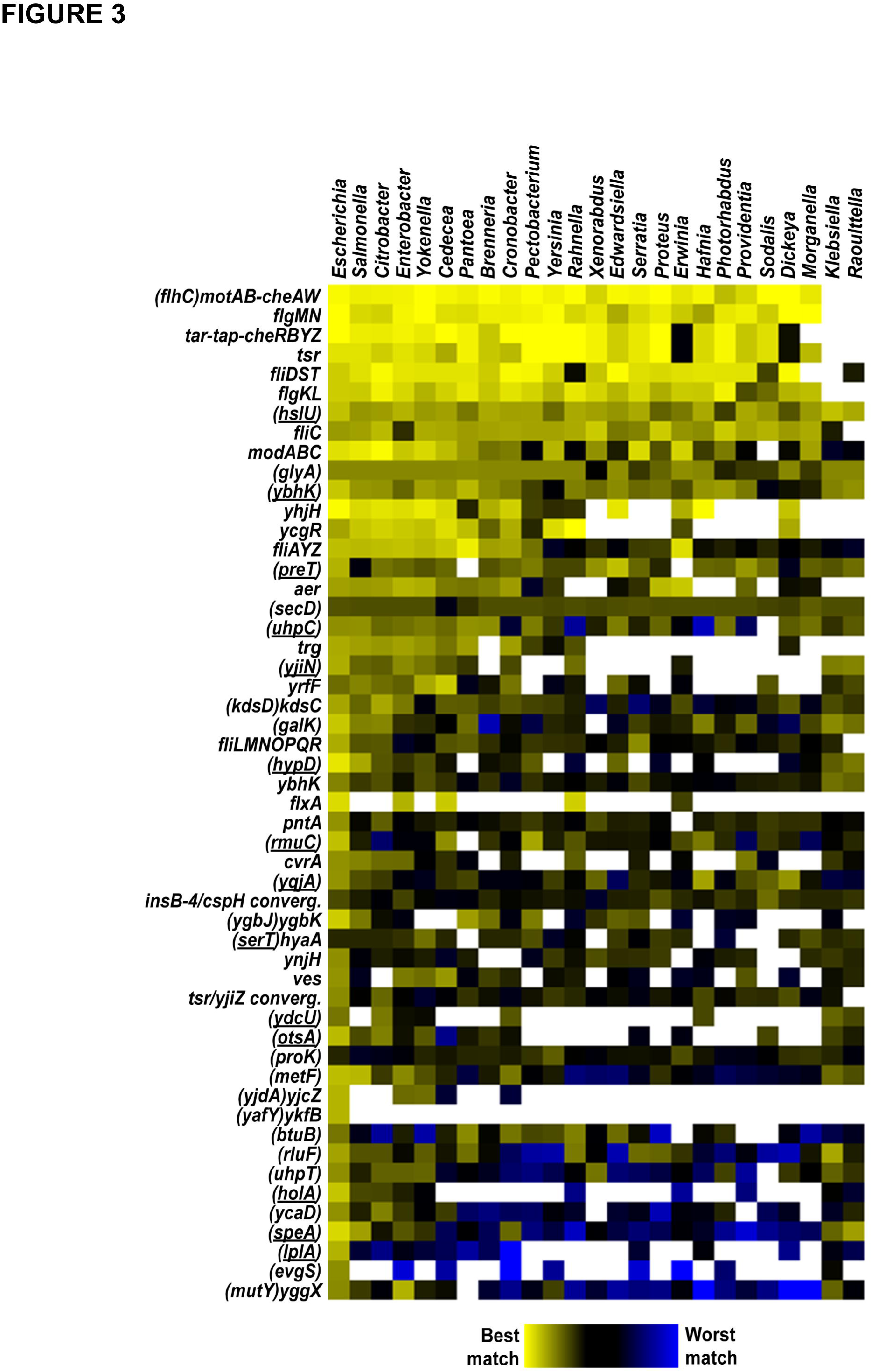
Sequence conservation of FliA binding sites between *E. coli* and related bacterial species. Heat-map depicting the match to the FliA consensus binding site for regions in the genomes of a range of bacterial species, where the region analyzed is homologous to a region surrounding a FliA binding site in *E. coli*. Genera are listed across the top. *E. coli* genes associated with the binding sites are listed to the left of the heat-map. Genes containing FliA binding sites are listed in parentheses. Underlining indicates that the putative promoter is in the antisense orientation relative to the overlapping gene. Genes not in parentheses are downstream of the corresponding FliA binding site. The color scale indicating the strength of the sequence match is shown below the heat-map. Empty squares in the heat-map indicate that the corresponding genomic region in *E. coli* is not sufficiently conserved in the species being analyzed.

### *Genome-wide mapping of the* Salmonella *Typhimurium FliA regulon*

*Salmonella enterica* and *E. coli* diverged approximately 100 million years ago and exhibit substantial drift at wobble positions [30]. As an independent, empirical test of FliA binding site conservation, we determined the genome-wide binding profile of *S enterica* serovar Typhimurium FliA using ChIP-seq of a C-terminally tagged derivative expressed from its native locus. A total of 23 high-confidence FliA binding sites were identified (Table 2, Figure 4A). Of these 23 sites, three are inside genes but within 300 bp of a gene start (13%; Figure 4B), and five are inside genes and far from a gene start (22%). No equivalent ChIP-seq peaks were identified using a control, untagged strain of *S.* Typhimurium. All 23 *S.* Typhimurium FliA binding sites are associated with a match to the consensus FliA motif (Figure 4C; MEME, E-value = 7.4e^−49^). As predicted by the sequence conservation analysis (Figure 3), FliA-dependent promoters upstream of key flagellar operons were conserved in *S.* Typhimurium. However, with the exception of the *motA* promoter that is located inside *flhC*, no intragenic FliA binding sites were found to be conserved between *E coli* and *S.* Typhimurium.

**Table 2.**
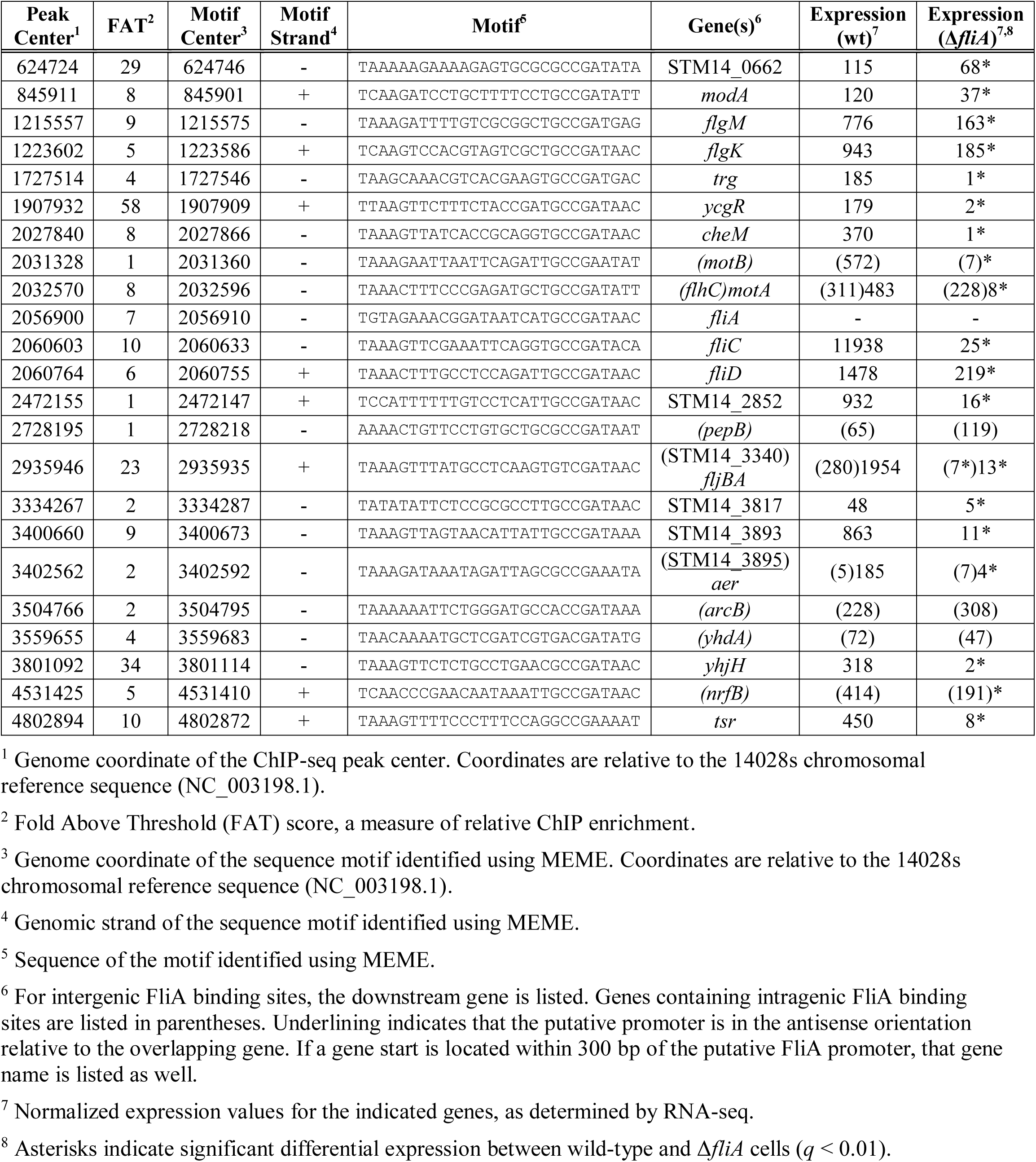
FliA regulon of *Salmonella* Typhimurium 14028s.

**Figure 4.**
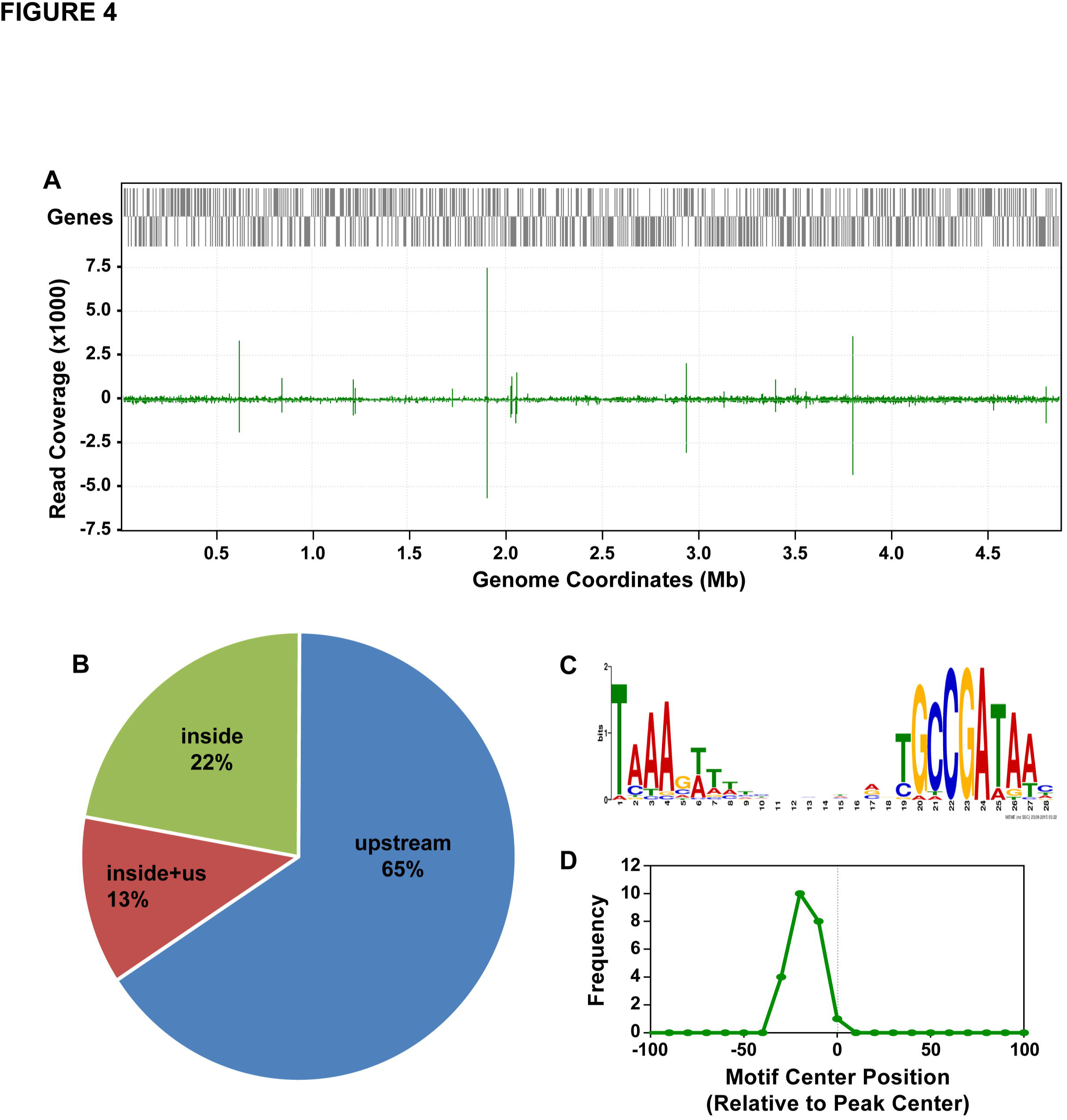
Identification of FliA binding sites in *Salmonella* Typhimurium using ChIP-seq. **(A)** Sequence read coverage across the *S.* Typhiumurium genome for a FliA ChIP-seq dataset. Annotated genes are indicated by gray bars. The green graph shows relative sequence read coverage, with “spikes” corresponding to sites of FliA association. **(B)** Pie-chart showing the distribution of identified FliA binding sites relative to genes. “Inside” = FliA binding within a gene. “Upstream” = FliA binding upstream of a gene. “Inside + us” = FliA binding within a gene but within 300 bp of a downstream gene start. **(C)** Enriched sequence motif associated with FliA binding sites identified by ChIP-seq. **(D)** Distribution of motifs relative to ChIP-seq peak centers for all FliA binding sites identified by ChIP-seq. Motifs are enriched in the region ~25 bp upstream of the peak center, relative to the motif orientation.

RNA-seq was used to assess FliA-dependent changes in gene expression by comparing wild-type and *ΔfliA* strains *of S.* Typhimurium (Figure 5). Overall, 344 genes were significantly differentially expressed between the two strains (*q*-value ≤ 0.01, fold-change ≥ 2), of which 36 were downstream of FliA binding sites identified by ChIP-seq (Table 2). The intragenic FliA binding sites within *flhC, STM14_3340*, and *STM14_3895* were associated with FliA-dependent regulation of the downstream genes, all of which are known flagellar genes. The other intragenic binding sites were not associated with detectable transcripts.

**Figure 5.**
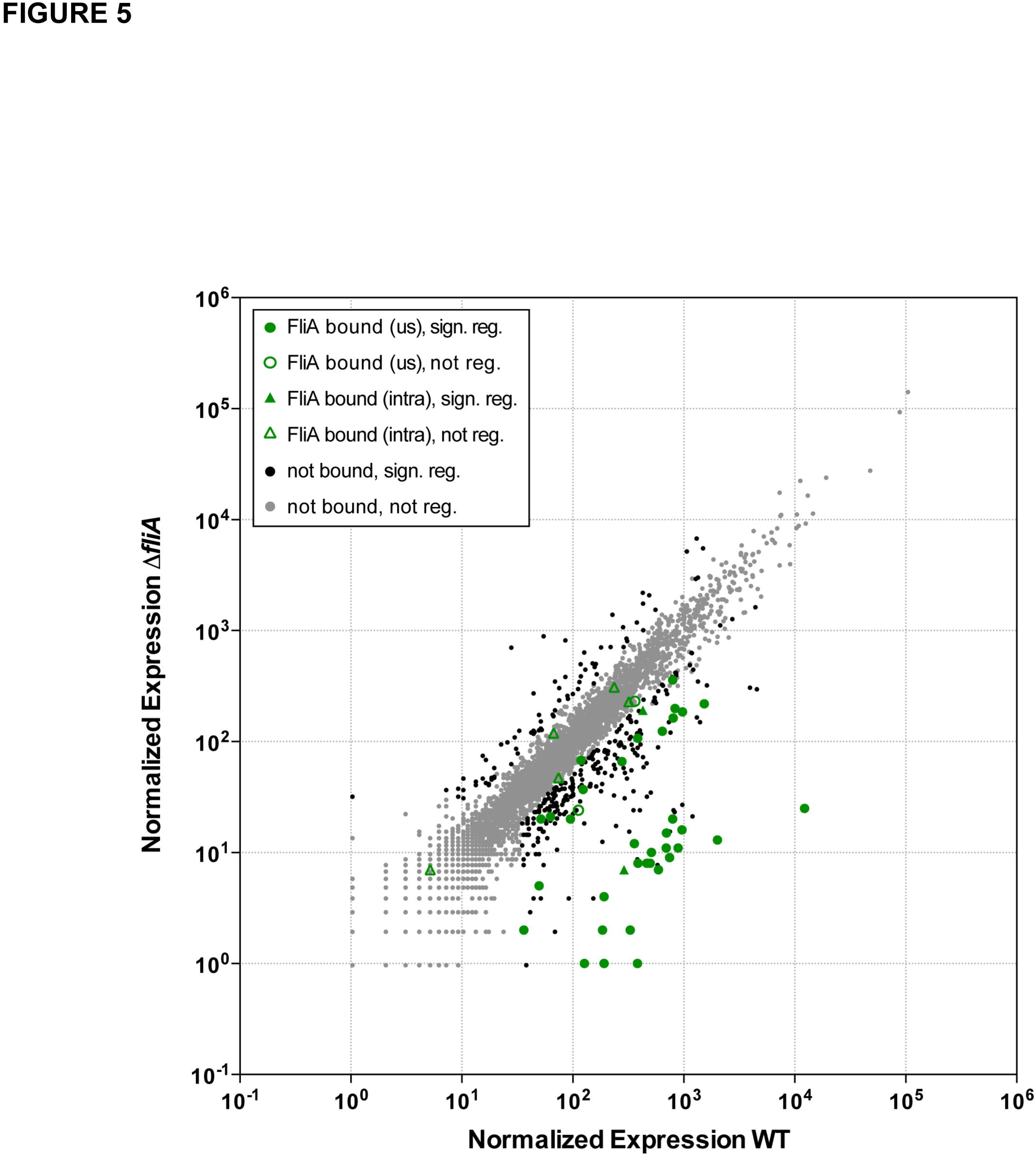
Transcriptome analysis of the FliA regulon in *Salmonella* Typhimurium. The scatter-plot shows normalized expression (see Methods) for each gene in *S.* Typhimurium for wild-type cells (14028s; x-axis) or Δ*fliA* cells (DMF088; y-axis). Gray dots represent genes that are not associated with a FliA binding site and are not significantly differentially expressed between wild-type and Δ*fliA* cells. Black dots represent genes that are not associated with a FliA binding site and are significantly differentially expressed between wild-type and Δ*fliA* cells. Green circles represent genes that are associated with an upstream FliA binding site. Green triangles represent genes that are associated with an internal binding site. Filled green circles/triangles indicate genes that are significantly differentially expressed between wild-type and *ΔfliA* cells. Empty green circles/triangles represent genes that are not differentially expressed between wild-type and *ΔfliA* cells.

### The motA promoter within flhC constrains evolution of the FlhC protein

Although most intragenic FliA promoters in *E. coli* are not well conserved in other species, the *motA* promoter, located inside *flhC*, is highly conserved (Figure 3). However, it is unclear whether this conservation is due to selective pressure on the promoter or on the amino acid sequence of FlhC, which is encoded by the same DNA. As expected given the conservation of the *motA* promoter inside *flhC*, the two FlhC amino acids, Ala177-Asp178, that are encoded by sequence overlapping the −10 region, are highly conserved among γ-proteobacteria (Figure 6A). Strikingly, the amino acids flanking the Ala-Asp sequence are poorly conserved (Figure 6A), leading us to hypothesize that the Ala-Asp motif is conserved due to selective pressure on the *motA* promoter, rather than on the amino acids themselves. To test this hypothesis, we determined whether Asp178 is required for FlhC function. We created a strain of motile *E. coli* MG1655 in which the *flhDC* promoter is transcriptionally active, but *flhC* is replaced with a cassette containing *thyA* under the control of a constitutive σ^70^ promoter. Thus, this strain lacks the *motA* promoter, but we reasoned that *motA* would be co-transcribed with *thyA* (Figure 6B). We then introduced either wild-type FlhC or D178A FlhC from a plasmid, or an empty vector control. Cells containing the empty vector control were non-motile, as expected given that they lack FlhC (Figure 6B). By contrast, cells expressing wild-type FlhC from the plasmid were fully motile. Strikingly, cells expressing D178A FlhC were also fully motile (mean motility level relative to wild-type FlhC of 0.97 ± s.d. 0.09, n = 3; Figure 6B). We conclude that the conserved Asp178 is not required for FlhC function.

**Figure 6.**
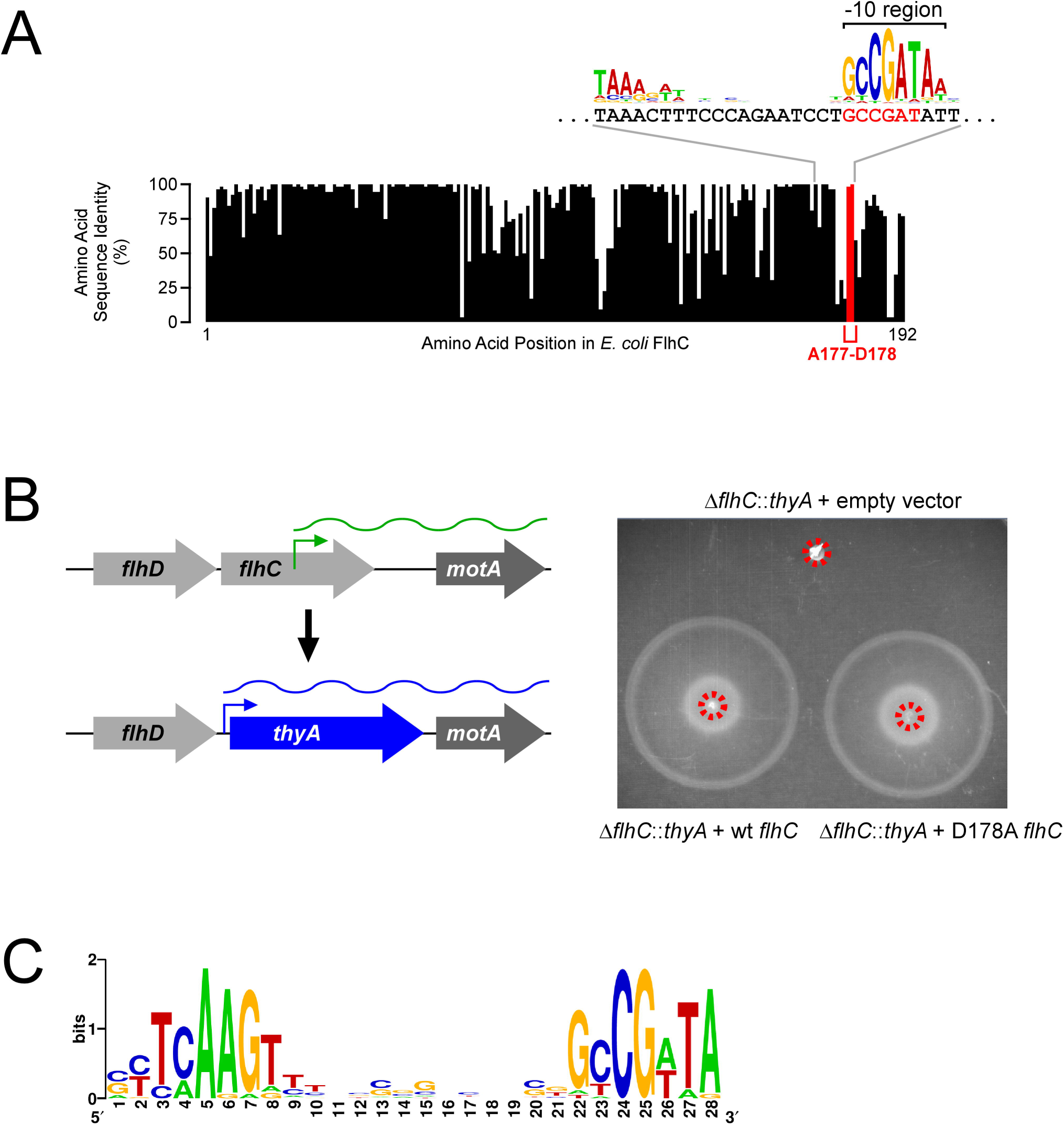
The FliA promoter within *flhC* constrains evolution of FlhC amino acid sequence. **(A)** Sequence conservation of FlhC amino acid sequence between *E. coli* and 51 other γ-proteobacterial species. The graph indicates the level of identity across all species analyzed for each amino acid in FlhC. Data for Ala177 and Asp178 are highlighted in red. The nucleotide sequence of *flhC* in the *motA* promoter region is indicated, aligned with the previously reported FliA binding motif logo [17]. Codons 177 and 178 are shown in red. **(B)** Motility assay for *ΔflhC*∷*thyA E. coli* (CDS105) containing either empty vector (pBAD30), or plasmid expressing wild-type FlhC (pCDS043) or D178A mutant FlhC (pCDS044). Red circles indicate the inoculation sites. Plates were incubated for 7 hours. The schematic to the left of the plate image shows how the strain was constructed. **(C)** Enriched sequence motif found in the *flhC-motA* intergenic region in species analyzed for which FlhC Asp178 is not conserved. This motif is a close match to the known FliA binding site consensus.

To further investigate the conservation of the Ala-Asp motif in FlhC, we aligned the sequences of FlhC homologues from 99 different proteobacterial species, each from a different genus, in which *motA* is positioned immediately downstream of *flhC.* Although Ala177 and Asp178 are well conserved across these species (conserved in 70% and 56% of species, respectively), we identified 44 species in which Asp178 is not conserved. We reasoned that if Asp178 is broadly conserved due to selective pressure on the overlapping *motA* promoter, species in which Asp178 is not conserved are likely to have repositioned the *motA* promoter. To test this hypothesis, we extracted the intergenic sequences between *flhC* and *motA* for each of the 44 species where Asp178 is not conserved. Consistent with our hypothesis, we identified a strongly enriched sequence motif (n = 19; E-value = 1.5e^−32^) corresponding to a consensus FliA promoter (Figure 6C). By contrast, when we repeated this analysis for the 55 species where Asp178 is conserved, we did not observe enrichment of a FliA promoter motif in the *flhC-motA* intergenic region. We conclude that the selective pressure on Asp178 is lost in species that reposition the *motA* promoter to the *flhC*-*motA* intergenic region.

## DISCUSSION

### Most FliA Binding Sites are Active Promoters for Unstable RNAs

Most FliA binding sites identified by ChIP-seq display FliA-dependent promoter activity when fused upstream of the *lacZ* reporter gene (Figure 1). Many of these FliA binding sites, and some additional sites that had inactive *lacZ* fusions, are associated with correctly positioned TSSs and NET-seq signal from published studies [26,27]. Together, these data suggest that almost all FliA binding sites represent transcriptionally active FliA-dependent promoters, regardless of their location relative to protein-coding genes. The small subset of FliA binding sites that appear to be transcriptionally inert were amongst the most weakly bound sites detected by ChIP-seq [17]. Three of these sites have at least one mismatch to key −10 region residues [22], suggesting that the sites are unlikely to be active promoters, or are so weakly transcribed that their activity is undetectable using standard assays.

Although most intragenic FliA binding sites are likely to represent active promoters, they are not associated with the transcription of stable RNAs, since we previously detect very few such RNAs using standard RNA-seq [17]. We conclude that most intragenic FliA promoters drive transcription of unstable RNAs. This is consistent with the previously described phenomenon of “pervasive transcription” that generates large numbers of short, unstable transcripts, primarily from promoters within genes [1,2]. Intragenic promoters typically drive transcription of non-coding RNAs. Transcription of these RNAs is rapidly terminated by Rho [31], and the transcripts are rapidly degraded by RNases [1,2].

### Limited conservation of the FliA regulon outside of core flagellar genes

DNA sequences conferring beneficial functions are more likely to be evolutionarily conserved, while neutral or detrimental sequences are lost by drift or negative selection, respectively. As expected, most flagella-associated FliA promoters are highly conserved at the sequence level (Figures 3 + 4). Of the intragenic FliA binding sites, only those that drive transcription of an mRNA for a downstream gene appear to be at all functionally conserved. A few intragenic promoters, such as those within *hslU, glyA*, and *ybhK*, are conserved at the sequence level between *E. coli* and many species. However, the fact that these sites are also conserved in two genera not encoding *fliA, Klebsiella* and *Raoultella*, suggests that the DNA sequences are maintained for reasons independent of FliA, most likely positive selection on the codons for the overlapping protein-coding genes.

To experimentally validate the sequence-based conservation predictions, we performed ChIP-seq on *S.* Typhimurium FliA. As predicted based on sequence conservation, all key flagellar promoters were functionally conserved, except the one upstream of *fliLMNOPQR.* In *E. coli*, this operon is primarily transcribed from a σ^70^ promoter that is activated by FlhDC [17,24,25]. Conservation of the σ^70^ promoter and FlhDC regulation would ensure that these genes are coordinately regulated with other flagellar genes in *S.* Typhimurium, potentially relieving the selective pressure to maintain the FliA promoter. Our ChIP-seq data indicate the only intragenic FliA promoter functionally conserved between *E. coli* and *S.* Typhimurium is that within *flhC.* While specific intragenic FliA binding sites were not conserved, *S.* Typhimurium FliA binds multiple intragenic sites. This suggests that the factors affecting FliA specificity, or lack thereof, are similar between *E. coli* and *S.* Typhimurium, and that the phenomenon of intragenic FliA promoters is conserved even if the specific promoters are not.

It should be noted that lack of conservation of specific promoters does not necessarily indicate a lack of functional importance, but could instead reflect lineage-specific evolution. Indeed, regulatory small RNAs are often poorly conserved, even between closely related species [32-34]. Consistent with this, one of the two stable, FliA-transcribed non-coding RNAs - that transcribed from within *uhpT -* is likely a functional regulator, despite a lack of promoter conservation. A recent study detected numerous Hfq-mediated interactions between mRNAs and RNA originating from the 3’ end of *uhpT* [35]. Although the *uhpT* sequences from these interactions map to locations downstream of the sRNA predicted by RNA-seq [17], an earlier microarray study and NET-seq data suggest that the FliA-transcribed sRNA extends further downstream [27,36]. The other stable, FliA-transcribed non-coding RNA - that transcribed from within *hypD -* was not detected in any sRNA:mRNA interactions [35], suggesting that it is not functional. Unstable, FliA-transcribed non-coding RNAs are also likely not functional, given their transient nature, and the lack of promoter conservation.

Intragenic FliA promoters likely arise as a result of sequence drift during evolution. The prevalence of intragenic FliA promoters in *E. coli* and *S*. Typhimurium suggests that they do not substantially impact expression of the overlapping genes. Consistent with this, we detected significant FliA-dependent regulation of only three *S.* Typhimurium genes with an internal FliA site (Figure 5; Table 2); one of these genes is immediately upstream of a FliA-transcribed flagellar gene, and another is also a downstream gene in a FliA-transcribed operon. While most intragenic FliA promoters are unlikely to be individually functional, the phenomenon of widespread intragenic FliA sites may be functional. For example, intragenic FliA sites could titrate cellular FliA, thereby sensitizing other FliA promoters to the level of FliA expression [37]. Alternatively, titration of FliA could reduce stochasticity in effective FliA levels, by requiring that FliA levels be maintained at higher levels. These functions would be independent of the specific locations of FliA promoters, and more dependent on the number and strength of promoters. Spontaneous creation of FliA binding sites by genetic drift may also provide a source of novel, functional FliA promoters, e.g. if there is a selective advantage of coordinately regulating the downstream gene with flagellar genes.

### *The* motA *promoter inside* flhC *constrains the evolution of FlhC*

Although most intragenic FliA promoters are not conserved, the promoter within *flhC* is the most highly conserved of all FliA promoters. This promoter has been described previously, and drives transcription of the *motAB*-*cheAW* operon mRNA [17,38,39]. FliA promoters require a stringent match to the consensus promoter sequence [22], and this is reflected by the high information content in the sequence motif associated with FliA binding, especially in the −10 region (Figure 4C) [17]. Hence, conservation of an intragenic FliA promoter is likely to result in conservation of the amino acid sequence for the overlapping codons. The −10 region of the FliA promoter in *flhC* corresponds to an Ala-Asp motif in the FlhC protein. This motif is broadly conserved. Multiple independent lines of evidence support the idea that the Ala-Asp sequence motif is conserved due to selective pressure on the intragenic FliA promoter and not on the amino acids themselves: (i) amino acids close to the Ala-Asp motif that are not associated with FliA promoter elements are poorly conserved (Figure 6A); (ii) the Ala-Asp motif is not present in the X-ray crystal structure of FlhDC [40], suggesting that it is in a disordered region; (iii) Asp178 does not detectably contribute to FlhC function (Figure 6B); and (iv) in proteobacterial species where *flhC* and *motA* are adjacent genes but FlhC Asp178 is not conserved, an alternative FliA promoter is often located in the intergenic region between *flhC* and *motA* (Figure 6C). Thus, even in cases where the specific FliA promoter inside *flhC* is not conserved, the presence of a FliA promoter upstream of *motA* is conserved. If the FliA promoter inside *flhC* were conserved because of selective pressure on the Ala-Asp motif, we would expect that (i) surrounding amino acids would also be conserved, regardless of whether they are encoded in sequence overlapping key FliA promoter elements, (ii) the Ala-Asp motif would be part of an important structural motif, (iii) Asp178 would be required for motility, and (iv) in species where Asp178 is not conserved, there would be no selective pressure to acquire an alternative FliA promoter for *motA*. We therefore conclude that the amino acid sequence of FlhC is constrained by the internal promoter for *motA*. Thus, the evolution of FlhC protein sequence is directly impacted by the function of the downstream gene.

### The potential for an abundance of bacterial duons

A recent study [15] coined the term “duon” to refer to sequences that are under selective pressure for two distinct functions, and identified large numbers of potential duons in the human genome. While the specific findings of that study have been questioned [41], the FliA promoter inside *flhC* clearly represents a bacterial duon. We propose that duons are likely to occur far more frequently in bacteria than in eukaryotes. The compact nature of bacterial genomes causes them to be gene-dense, greatly limiting the non-coding sequence space; in *E. coli*, ~90% of the genome is protein-coding, in stark contrast to the human genome, which is <2% protein-coding. Consistent with the paucity of non-coding sequence in bacterial genomes, numerous intragenic binding sites have been identified for transcription factors and σ factors [2,4–11]. In some cases, low stringency in the DNA sequence requirements for binding may limit the potential for duons. For example, there are many intragenic σ^70^ promoters in *E. coli* [3], but σ^70^ promoters can still be active with multiple mismatches to the consensus [3]. Hence, even if an intragenic σ^70^ promoter is under positive selective pressure, it could acquire mutations that alter the overlapping coding potential without affecting promoter strength. However, bacterial transcription factors tend to have high information content binding sites, especially compared to their eukaryotic equivalents [42,43]. This suggests that functional conservation of intragenic transcription/σ factor binding sites in bacteria will often constrain evolution of the overlapping gene.

Identification of duons requires further investigation of intragenic regulatory sites. Although numerous intragenic binding sites have been identified, their regulatory capacity is often unclear, and their conservation has not been analyzed. Nonetheless, there are described examples of intragenic σ factor binding that likely constitute duons. First, an intragenic promoter for the alternative σ factor, σ^24^, is conserved both at the sequence level and functionally [44,45]. This promoter drives transcription of a non-coding, regulatory RNA, MicL, that is also conserved [44]. Hence, both the promoter and non-coding RNA might represent dual-usage sequence. Second, an alternative σ factor, σ^54^, binds many intragenic sites in *E. coli* and *S.* Typhimurium that are conserved both at the sequence level and functionally [4,46], suggesting that they may be duons. Since conserved intragenic σ^54^ binding sites are likely to be promoters for downstream genes [4], evolution of the amino acid sequence of proteins encoded by genes containing σ^54^ promoters may often be constrained by the function of the downstream gene.

Extrapolating from our data for FliA, the majority of intragenic transcription/σ factor binding sites are likely to be non-functional, and hence not under positive selective pressure. These sites would therefore not be duons. Even though the complete regulons of most *E. coli* transcription/σ factors remain to be mapped, thousands of intragenic sites have already been identified, implying that there are thousands more sites yet to be discovered. Even if only a small fraction of intragenic sites are under positive selective pressure, this would indicate the existence of many duons. Hence, our data suggest that the evolutionary impact of intragenic regulatory sequences should be considered more broadly, as it is likely to be an important factor shaping bacterial genome evolution.

## MATERIALS AND METHODS

### Strains, plasmids, and growth conditions

All bacterial strains and plasmids used in this study are listed in Table 3. All oligonucleotides used in this study are listed in Table S1. All *E. coli* strains are derivatives of the motile MG1655 strain (DMF36) described previously [17]. To construct strains used for β-galactosidase assays, the native *lacZ* gene of DMF36, or the isogenic Δ*fliA* strain (DMF40) [17] was replaced by *thyA* using FRUIT recombineering [47] with oligonucleotides JW5397 and JW5398, generating strains DMF122 and DMF123, respectively. *flhC* and 106 bp downstream sequence was replaced with *thyA* in DMF36 using FRUIT recombineering [47] to generate strain CDS105. *Salmonella* strains are derivatives of *S enterica* serovar Typhimurium 14028s [48]. *S.* Typhimurium FliA was N-terminally epitope tagged with a 3x-FLAG tag at the native chromosomal locus using FRUIT recombineering [47], generating strain DMF087. The *S.* Typhimurium Δ*fliA* strain, DMF088, was constructed using FRUIT recombineering [47].

**Table 3.**
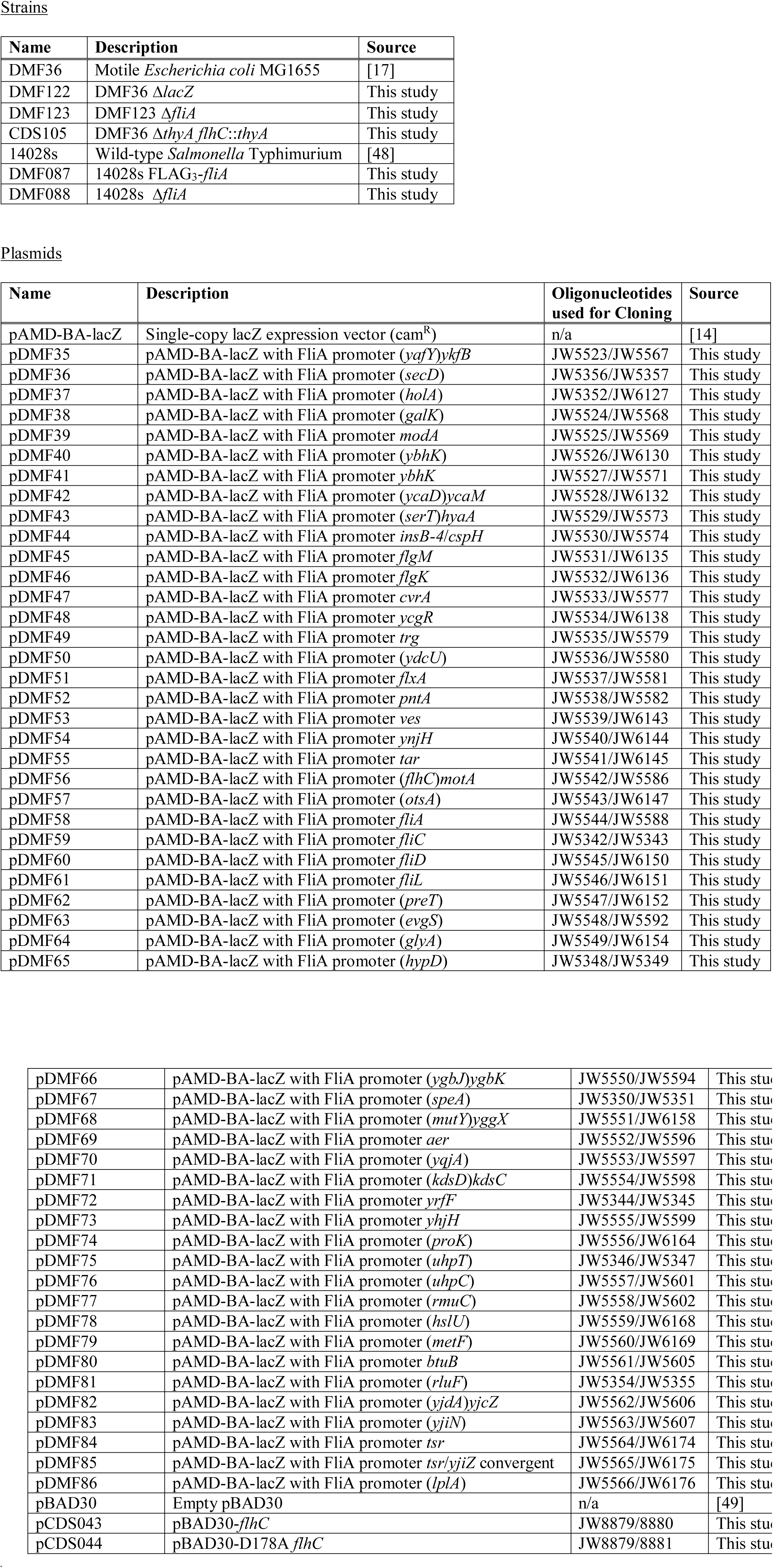
Strains and Plasmids used in this study.

Wild-type *flhC* was PCR amplified using oligonucleotides JW8879 and JW8880, and cloned into the *Sac* I and *Sal* I restriction sites of pBAD30 [49] using the In-Fusion method (Clontech) to generate pCDS043. D178A mutant *flhC* was PCR amplified using oligonucleotides JW8879 and JW8881, and cloned as described for wild-type *fhlC*, to generate pCDS044. Transcriptional fusions of putative FliA promoters to *lacZ* were constructed in plasmid pAMD-BA-lacZ [14]. Putative promoter regions (nucleotide positions −200 to +10, relative to the predicted TSS) were PCR-amplified from MG1655 cells. PCR products were cloned into pAMD-BA-lacZ cut with *SphI* and *NheI* using the In-Fusion method (Clontech). Oligonucleotides used for the plasmid cloning are listed in Table 3.

For all experiments, subcultures were grown in LB at 37 °C, with aeration, to OD_600_ 0.5-0.7.

### Construction of promoter-lacZ fusions, and β-galactosidase assays

Transcriptional *lacZ* promoter fusion plasmids were transformed into Δ*lacZ* strains with (DMF122) or without *fliA* (DMF123). Promoter activity was assessed by β-galactosidase assay, as previously described [14].

### Analysis of published TSS data

To determine whether FliA binding sites were associated with TSSs, the published list of TSS locations derived from differential RNA-seq was used [26]. Orientation of putative FliA promoters was determined based on associated motifs. For each putative FliA promoter, the distance from the motif center to each downstream TSS on the correct strand was calculated. All pairwise distances <500 bp are plotted in Figure 2A. As a control, a randomized TSS dataset was generated with the same total number and distribution (with respect to strand and gene location) as the experimental dataset. The analysis was repeated with this dataset.

### Analysis of published NET-seq data

Raw sequencing data files from NET-seq experiments [27] were obtained and mapped to the *E. coli* MG1655 genome using CLC Genomics Workbench. Sequence read depths at positions surrounding putative FliA promoters were calculated using a custom Python script. For FliA binding sites associated with a TSS, the NET-seq read coverage was calculated at every position from −100 to +100 relative to the TSS. For FliA binding sites not associated with a TSS, a TSS was predicted to be located 20 bp downstream of the motif center, and NET-seq read coverage was calculated from −100 to +100 relative to this position. For each region, NET-seq read coverage was normalized to local minimum and maximum values. Normalized read coverage was plotted as a heat map in Figure 3B.

### FliA binding site conservation analysis

The locations of all *E. coli* FliA binding sites described previously [17] were used to identify homologous sequences in 24 other species, as described previously [4]. These sequences were scored for matches to a Position Specific Scoring Matrix derived from the identified FliA binding sites in *E. coli* [17], as described previously [4].

### ChIP-seq of S. Typhimurium FliA

ChIP-seq was performed with strains DMF087 (FliA-FLAG3) or 14028s (untagged control) as previously described [14]. Sequence reads were mapped to the *S*. Typhimurium 14028s genome using CLC Genomics Workbench (Version 8). Peaks were called using a previously described analysis pipeline [17]. Three peaks with a FAT score of 1 were identified in the control dataset; these peaks were all >30 kbp from any putative FliA binding site.

### RNA-seq

RNA-seq was performed with strains 14028s and DMF088 as previously described [14]. Read mapping and differential expression analysis were performed with Rockhopper [50]. The normalized expression values and indicators of statistical significance in Table 2 were generated using Rockhopper.

### Analysis of FlhC sequence conservation

We used the RSAT “Comparative Genomics/Get Orthologs” tool (default parameters except we required 50% amino acid sequence identity; [51]) to identify 52 FlhC homologues from γ-proteobacterial species, each from a different genus. We aligned protein sequences using MUSCLE (v3.8, default parameters; [52]), and for each FlhC homologue we counted matches at each amino acid position to the aligned *E. coli* FlhC sequence.

### Identification of enriched sequence motifs in flhC-motA intergenic regions

We used the RSAT “Comparative Genomics/Get Orthologs” tool (default parameters except required 40% amino acid sequence identity; [51]) to identify 131 FlhC homologues from proteobacterial species, each from a different genus. To determine whether the *flhC* and *motA* genes are adjacent in each of the 131 species selected, we first used the RSAT “Comparative Genomics/Get Orthologs” tool (default parameters except required 40% amino acid sequence identity; [51]) to extract 100 bp of sequence immediately downstream of *flhC* for each species. We then searched for open reading frames similar to that of *motA* using BLASTX (v2.2.3, hosted on EcoGene 3.0, default parameters, searching against the *E. coli* annotated proteome; [53,54]). We discarded 32 FlhC sequences for which there was no BLASTX match to MotA with the corresponding sequence downstream of *flhC*. We aligned the remaining 99 protein sequences using MUSCLE (v3.8, default parameters; [52]), and for each FlhC homologue we counted matches at each amino acid position to the aligned *E. coli* FlhC sequence.

We used the RSAT “Comparative Genomics/Get Orthologs” tool [51] to extract intergenic sequence downstream of *flhC* for the 99 FlhC homologues from genomes where *flhC* and *motA* are adjacent genes. We discarded intergenic sequences <50 bp. We used MEME (v4.12.0, default settings except “look on given strand only”; [55]) to identify enriched sequence motifs in intergenic regions from species where FlhC Asp178 is conserved or is not conserved, respectively.

### Motility assays

Motility assays were performed as previously described [17].

## ACCESSION NUMBERS

Raw ChIP-seq and RNA-seq data are available from the EBI ArrayExpress repository using accession numbers E-MTAB-6048 (RNA-seq) and E-MTAB-6049 (ChIP-seq).

## ACKNOWLEDGEMENTS

We thank the Applied Genomic Technologies Core Facility for Sanger sequencing, the University at Buffalo Next Generation Sequencing Core Facility for Illumina sequencing, and the Wadsworth Center Media and Glassware Core facilities for media and glassware. We thank David Grainger, Keith Derbyshire, and members of the Wade group for helpful discussions.

